# Association between the extent of DNA methylation at the CpG sites of *HIF3A* and parameters of obesity in the general Japanese population

**DOI:** 10.1101/2020.11.30.403709

**Authors:** Genki Mizuno, Hiroya Yamada, Eiji Munetsuna, Mirai Yamazaki, Yoshitaka Ando, Ryosuke Fujii, Yoshiki Tsuboi, Atsushi Teshigawara, Itsuki Kageyama, Keisuke Osakabe, Keiko Sugimoto, Hiroaki Ishikawa, Naohiro Ichino, Yoshiji Ohta, Koji Ohashi, Shuji Hashimoto, Koji Suzuki

**Affiliations:** Department of Preventive Medical Sciences, Fujita Health University School of Medical Sciences, Toyoake, Aichi, Japan; Department of Joint Research Laboratory of Clinical Medicine, Fujita Health University Hospital, Toyoake, Aichi, Japan; Department of Hygiene, Fujita Health University School of Medicine, Toyoake, Aichi, Japan; Department of Biochemistry, Fujita Health University School of Medicine, Toyoake, Aichi, Japan; Department of Medical Technology, Kagawa Prefectural University of Health Sciences, Takamatsu, Kagawa, Japan; Department of Clinical Biochemistry, Fujita Health University School of Medical Sciences, Toyoake, Aichi, Japan; Department of Clinical Physiology, Fujita Health University School of Medical Sciences, Toyoake, Aichi, Japan; Department of Chemistry, Fujita Health University School of Medicine, Toyoake, Aichi, Japan

## Abstract

Obesity is a major public health problem worldwide owing to the substantial increase in risk of metabolic diseases. Hypoxia-inducible factors (HIFs) regulate transcriptional responses to hypoxic stress. DNA methylation in the CpG sites of intron 1 of *HIF3A* is associated with body mass index in the whole blood and adipose tissue. This study investigates the correlation between DNA methylation of *HIF3A* and parameters of obesity, including thickness of visceral (VAT) and subcutaneous adipose tissues, in the general Japanese population. Participants (220 men and 253 women) who underwent medical examination were enrolled in this cross-sectional study. We used pyrosequencing to quantify DNA methylation (CpG sites of cg16672562, cg22891070, and cg27146050) in *HIF3A*. DNA methylation of *HIF3A* was only different in women. Multiple regression analysis showed that DNA methylation level at cg27146050 was associated with thickness of VAT in women. DNA methylation level at cg27146050 also correlated with body mass index and percentage of body fat in women after excluding smokers and non-smokers who quit smoking with the last 5 years. DNA methylation in the CpG site (cg27146050) of *HIF3A* correlated with parameters of obesity in Japanese women.

## Introduction

Obesity is a major public health concern worldwide. In obese individuals, non-esterified fatty acids, adipokines, and other factors are extensively released from adipose tissues, thereby leading to abnormalities in obesity-related cell functions^1^. Consequently, obesity induces various diseases, such as insulin resistance, type 2 diabetes, and cardiovascular disease^2, 3^. Thus, obesity is a risk factor for various metabolic diseases, and preventing obesity results in the prevention of metabolic diseases. Recent years have seen the diversification of lifestyle and eating habits that have increased the number of obese individuals globally^4^. Lifestyle, environmental factors, and genetic factors trigger obesity^5, 6^. Lifestyle and/or environmental factors cause epigenetic alterations in several health conditions, such as obesity and metabolic disease^7–10^.

DNA methylation is an epigenetic mechanism that regulates gene expression by adding a methyl donor to cytosine to enable the regulation of transcription^11^. Lifestyle factors, including dietary habits, modulate DNA methylation^12^. Several animals^13–15^ and epidemiological studies^16–18^ have shown that environmental factors, including food intake, tobacco smoking, and alcohol consumption, cause DNA methylation in the blood or tissues. Moreover, global DNA hypermethylation in leukocytes is associated with increased risk of cardiovascular diseases in the general Japanese population^19^. Thus, DNA methylation may be a novel biomarker for metabolic diseases caused by environmental factors and lifestyles.

Dick et al.^20^ conducted two epigenetic genome-wide analyses to show the increase in DNA methylation at three CpG sites (cg16672562, cg22891070, and cg27146050) in intron 1 of *HIF3A* in the blood was associated with body mass index (BMI). Similarly, Main et al.^21^ and Wang et al.^22^ demonstrated that DNA methylation in *HIF3A* in the blood is associated with BMI in patients with type 2 diabetes and childhood obesity, respectively. Isoforms of HIF are constitutively expressed in mammalian cells and regulate transcriptional response to hypoxic stress^23, 24^. HIFs are unstable at normal oxygen levels in mammalian cells. The reduction in normal cellular oxygen levels caused by environmental factors, diseases, effusion of blood, and adiposity stabilize HIFs, thereby enabling its nucleocytoplasmic translocation and binding to the hypoxia response element in the promoter of target genes and regulating target gene transcription and expression. Pfeiffer et al.^25^ have shown that methylation of *HIF3A* in the adipose tissue correlates with dysfunctional human subcutaneous adipose tissue (SAT) and visceral adipose tissue (VAT). These studies indicate that DNA methylation of *HIF3A* is associated with the development of obesity, and may be an obesity-related factor worldwide. Furthermore, DNA methylation of *HIF3A* in the blood is associated with insulin resistance in patients with type 2 or gestational diabetes^21, 26^. There are only a few reports on the association between DNA methylation of *HIF3A* and BMI in humans. The thickness of adipose tissues is a more reliable parameter of obesity as compared to BMI that is an indirect parameter. To the best of our knowledge, there is no study on the correlation between DNA methylation of *HIF3A* and thickness of adipose tissues, such as VAT and SAT, that directly reflects obesity.

In this study, we attempted to verify whether DNA methylation of *HIF3a* (CpG sites of cg16672562, cg22891070, and cg27146050) in the blood associated with the thickness of VAT and SAT in the general Japanese population. We further determined whether DNA methylation in *HIF3A* in the blood correlated with the thickness of VAT and SAT in Japanese non-smokers^27, 28^.

## Materials and methods

### Participants

This cross-sectional study was approved by the Ethics Review Committee of Fujita Health University (Approval number: HG19-069). We enrolled 473 participants (220 men and 253 women) who took part in the medical examination of the general (middle-aged) population in Yakumo town, Hokkaido, Japan, in August 2015^29, 30^. We obtained written informed consent from all the participants for the use of individual genome samples. Information on lifestyle habits was obtained from questionnaires.

### Measurements of obesity parameters

Parameters of obesity were measured as described previously^31^. Percentage of body fat (% body fat) was measured using bioelectrical impedance analysis with the Tanita MC780 multifrequency segmental body composition analyzer (Tokyo, Japan). The thicknesses of VAT and SAT were assessed using ultrasound with ProSound a7 and UST-9130 convex probe (Hitachi Aloka Medical, Ltd, Tokyo, Japan). Thickness of VAT and SAT were defined as the distance (cm) from the peritoneum to the vertebral bodies and depth (cm) from the skin to the linea alba, respectively.

### Blood test and determination of DNA methylation

Blood was collected during the medical examination of the general population, and the serum was separated from the blood by centrifugation at 2,000×g for 10 min at room temperature. For biochemical analysis of the blood, enzymes and components in the serum were assayed using an auto-analyzer (JCS-BM1650, Nihon Denshi Co., Tokyo, Japan) at Yakumo General Hospital.

DNA methylation was analyzed using the buffy coat obtained upon centrifugation of the blood collected in ethylenediaminetetraacetic acid (EDTA)-2Na-containing tubes under the same conditions as those used for blood biochemical tests. Genomic DNA was extracted from the buffy coat using the NucleoSpin Tissue kit (Takara, Shiga, Japan). Bisulfite conversion was performed using the Epitect Bisulfite Kit (Qiagen, Valencia, CA, USA). Polymerase chain reaction (PCR) was used to amplify the intron 1 of *HIF3A* using EpiTaqTM HS (for bisulfite-treated DNA; Takara, Shiga, Japan). Levels of DNA methylation were quantified using pyrosequencing with the PyroMark Q24 Advanced kit (Qiagen, Valencia, CA, USA) and analyzed using the parameters previously described^20–22^, including three CpG sites (Fig 1). Table 1 lists the sequences of primers used for PCR and pyrosequencing. The primers used for pyrosequencing were designed based on a previous study^25^ using PyroMark Assay Design 2.0 (Qiagen, Valencia, CA, USA).

**Fig. 1.**
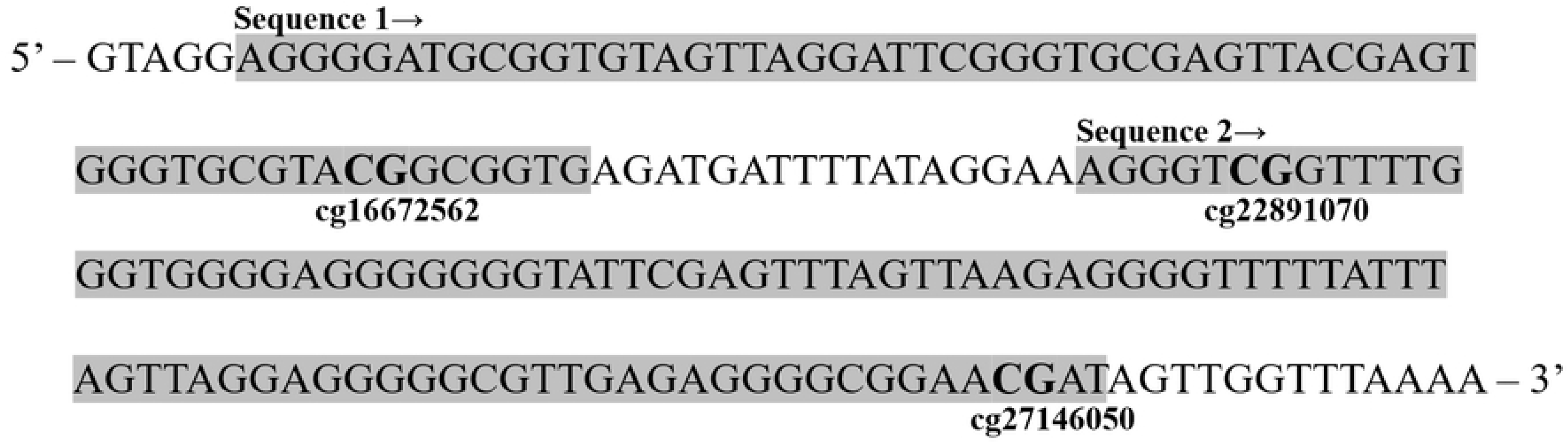
A target sequence of intron 1 region in *HIF3A* gene.

**Table 1.**
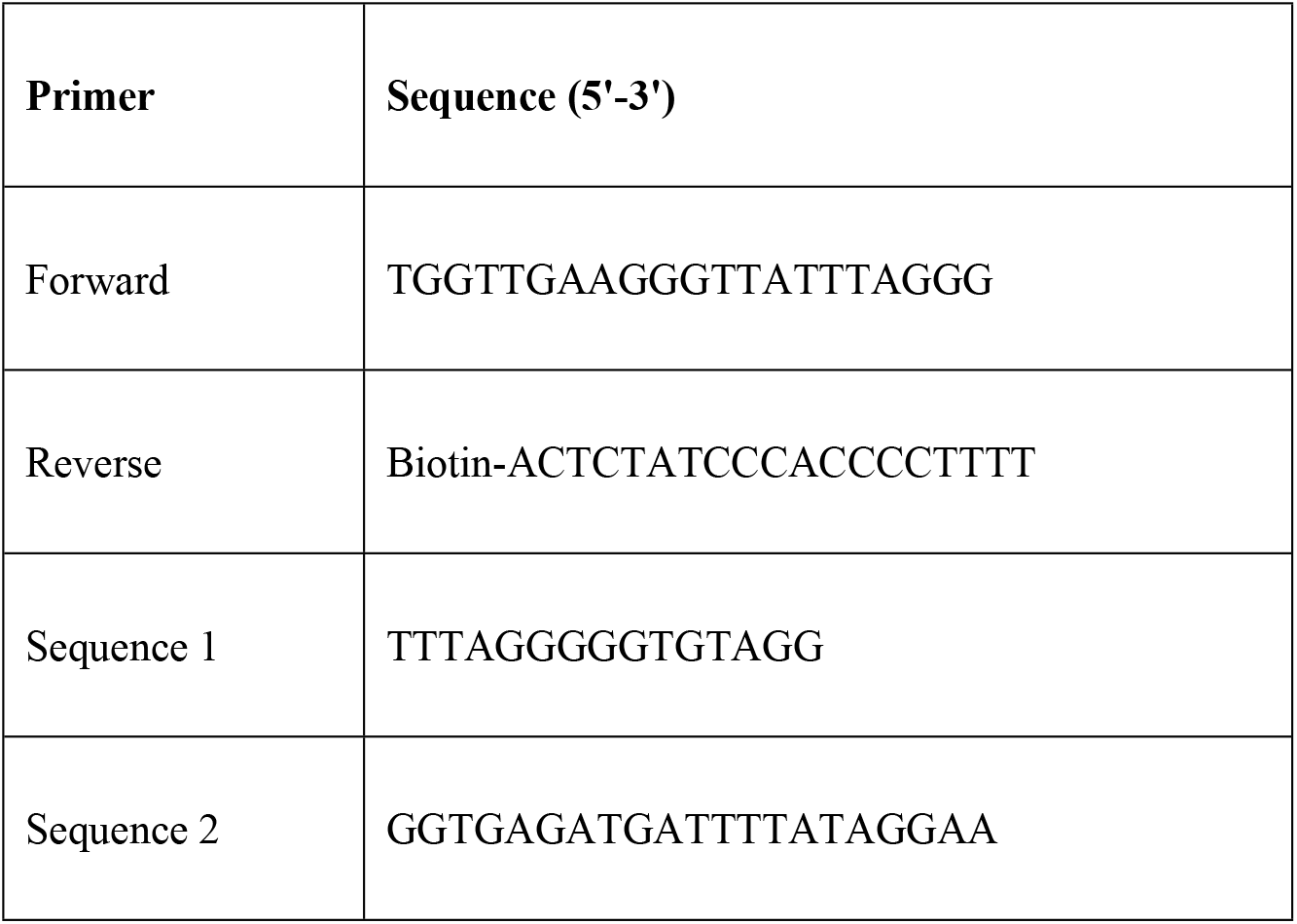
Sequences of primers used for PCR and Pyrosequence.

The target region of *HIF3A* gene DNA methylation analyzed by pyrosequence was decided based on previous studies. It has reported that the 3 CpG sites (cg16672562, cg22891070 and cg27146050) of intron 1 in *HIF3A* gene in the blood are associated with BMI in EWAS study.

### Statistical analysis

All statistical analyses were performed using JMP version 14.0 (SAS Institute, Cary, NC, USA). Serum aspartate transaminase (AST), alanine transaminase (ALT), triglyceride, and high-density lipoprotein (HDL) cholesterol levels have been represented by the geometric means and interquartile ranges owing to log-normal distribution. Other characteristics (including DNA methylation) have been represented as mean±standard deviation (SD). We analyzed the association of DNA methylation level at each CpG site of intron 1 in *HIF3A* with the parameters of obesity using single correlation and multiple linear regression and adjusted for age, systolic blood pressure, hemoglobin A1c, %neutrophil, smoking habit and exercise habit. For multiple testing, the Bonferroni method was used to counteract the problem of multiple comparisons. *P*<0.05 was considered statistically significant.

## Results

Table 2 lists the characteristics of the participants in this study. There were significant differences in various parameters of obesity between men and women, such as smoking habit and blood biochemical test, but not hemoglobin A1c and blood pressure. Moreover, DNA methylation levels at three CpG sites in intron 1 of *HIF3A* were significantly different between the sexes (Table 3).

**Table 2.** Characteristics of participants in this study.

|  | **Men** | **Women** | ***P*-value** |
| --- | --- | --- | --- |
| n | 220 | 253 |  |
| Age (years) | 66.3 ± 8.28 | 64.5 ± 8.00 | 0.017 <sup>a</sup> |
| Blood glucose (mg/dL) | 93.8 ± 14.5 | 87.6 ± 17.0 | <0.001 <sup>a</sup> |
| Hemoglobin A1c (%) | 5.80 ± 0.54 | 5.72 ± 0.55 | 0.190 |
| AST (IU/L) | 23.4 (19.0-26.8) | 21.6 (18.0-24.5) | 0.003 <sup>b</sup> |
| ALT (IU/L) | 23.1 (17.3-31.0) | 18.9 (14.0-24.0) | <0.001 <sup>b</sup> |
| Triglyceride (mg/dL) | 102.5 (72.0-144.8) | 86.3 (64.5-117.0) | <0.001 <sup>b</sup> |
| Total cholesterol (mg/dL) | 203.5 ± 32.4 | 217.5 ± 35.0 | <0.001 <sup>a</sup> |
| HDL cholesterol (mg/dL) | 52.4 (44.0-61.0) | 61.8 (53.0-72.0) | <0.001 <sup>b</sup> |
| LDL cholesterol (mg/dL) | 121.3 ± 30.2 | 127.0 ± 31.1 | 0.042 <sup>a</sup> |
| Systolic blood pressure (mmHg) | 134.8 ± 20.6 | 128.8 ± 19.2 | 0.001 <sup>a</sup> |
| Diastolic blood pressure (mmHg) | 79.8 ± 13.0 | 72.9 ± 12.6 | <0.001 <sup>a</sup> |
| Various parameters of obesity |  |  |  |
| BMI (kg/m<sup>2</sup>) | 24.1 ± 2.78 | 23.0 ± 3.60 | <0.001 <sup>a</sup> |
| VAT thickness (cm) | 64.6 ± 14.7 | 50.4 ± 11.7 | <0.001 <sup>a</sup> |
| SAT thickness (cm) | 14.3 ± 3.81 | 13.2 ± 4.62 | 0.006 <sup>a</sup> |
| % body fat | 23.7 ± 4.16 | 32.8 ± 6.13 | <0.001 <sup>a</sup> |
| Smoking habit, n (%) |  |  |  |
| Never | 45 (21) | 193 (77) | <0.001 <sup>c</sup> |
| Ever | 126 (57) | 40 (16) |  |
| Current | 49 (22) | 19 ( 7 ) |  |
| Exercise habit, n (%) |  |  |  |
| few | 102 (47) | 139 (55) | 0.366 |
| sometimes | 43 (20) | 44 (17) |  |
| 1 time/week | 24 (11) | 23 ( 9 ) |  |
| > 2 times/week | 49 (22) | 47 (19) |  |

Values are mean ± SD, geometric mean (25-75th parentheses), or n (%). *P* < 0.05 was considered
statistically significant. a: student t test, b: Wilcoxon test, c: Pearson's chi-square test

**Table 3.** DNA methylation levels (%) at *HIF3A* gene sites by pyrosequence analysis.

| CpG site | Men | Women | *P*-value |
| --- | --- | --- | --- |
| cg16672562 | 17.3 ± 5.12 | 20.1 ± 5.96 | <0.001 <sup>a</sup> |
| cg22891070 | 21.5 ± 7.37 | 24.9 ± 8.03 | <0.001 <sup>a</sup> |
| cg27146050 | 14.3 ± 4.74 | 17.1 ± 5.00 | <0.001 <sup>a</sup> |
| mean | 17.6 ± 5.00 | 20.6 ± 5.57 | <0.001 <sup>a</sup> |

Values are mean ± SD. *P* < 0.05 was considered statistically significant. a: student t test

Correlations between DNA methylation levels at the CpG sites in intron 1 of *HIF3A* and parameters of obesity were analyzed using single linear regression owing to the differences in DNA methylation of *HIF3A* in men and women (Tables 3 and 4). There was no significant correlation between DNA methylation level at each CpG site and the parameters of obesity in men and women.

**Table 4.** Single correlation analysis between *HIF3A* gene DNA methylation levels and obesity parameters in Japanese men and women.

| Men |  |  |  |  |  |  |  |  |
| --- | --- | --- | --- | --- | --- | --- | --- | --- |
| CpG site | BMI |  | VAT |  | SAT |  | % Body fat |  |
|  | *r* | *P* | *r* | *P* | *r* | *P* | *r* | *P* |
| cg16672562 | 0.087 | 0.210 | 0.082 | 0.244 | 0.029 | 0.682 | 0.074 | 0.295 |
| cg22891070 | 0.114 | 0.111 | 0.028 | 0.689 | 0.045 | 0.525 | 0.054 | 0.444 |
| cg27146050 | 0.013 | 0.854 | -0.006 | 0.937 | 0.050 | 0.478 | -0.021 | 0.766 |
| mean | 0.089 | 0.202 | 0.049 | 0.565 | 0.048 | 0.496 | 0.046 | 0.520 |
| Women |  |  |  |  |  |  |  |  |
| CpG site | BMI |  | VAT |  | SAT |  | % Body fat |  |
|  | *r* | *P* | *r* | *P* | *r* | *P* | *r* | *P* |
| cg16672562 | -0.029 | 0.661 | -0.066 | 0.317 | 0.025 | 0.704 | -0.026 | 0.690 |
| cg22891070 | 0.024 | 0.720 | -0.042 | 0.530 | 0.047 | 0.476 | 0.049 | 0.461 |
| cg27146050 | -0.038 | 0.562 | -0.109 | 0.099 | -0.028 | 0.679 | -0.045 | 0.497 |
| mean | 0.176 | 0.791 | 0.075 | 0.263 | 0.019 | 0.771 | 0.006 | 0.926 |

Table 5 shows the results of multiple linear regression analysis for the correlation between DNA methylation levels at CpG sites in intron 1 of *HIF3A* and the parameters of obesity. In men, there was no significant correlation between DNA methylation level at each CpG site and the parameters of obesity. In women, significant correlations were observed between DNA methylation level at cg27146050 and VAT thickness (*P*<0.05). However, DNA methylation levels at cg16672562 and cg22891070 did not significantly correlate with any parameter of obesity in women.

**Table 5.** Multiple linear regression analysis for correlations between *HIF3A* gene DNA methylation levels and obesity parameters.

| **Men** |  |  |  |  |  |  |  |  |
| --- | --- | --- | --- | --- | --- | --- | --- | --- |
| **CpG site** | **BMI** |  | **VAT** |  | **SAT** |  | **% Body fat** |  |
| | $\beta$ | $P$ | $\beta$ | $P$ | $\beta$ | $P$ | $\beta$ | $P$ |
| cg16672562 | 0.060 | 0.391 | 0.092 | 0.175 | -0.024 | 0.736 | 0.052 | 0.462 |
| cg22891070 | 0.081 | 0.241 | 0.030 | 0.656 | -0.007 | 0.922 | 0.040 | 0.570 |
| cg27146050 | -0.012 | 0.868 | 0.005 | 0.938 | -0.016 | 0.819 | -0.038 | 0.594 |
| mean | 0.059 | 0.402 | 0.050 | 0.467 | -0.016 | 0.818 | 0.027 | 0.706 |
| **Women** |  |  |  |  |  |  |  |  |
| **CpG site** | **BMI** |  | **VAT** |  | **SAT** |  | **% Body fat** |  |
| | $\beta$ | $P$ | $\beta$ | $P$ | $\beta$ | $P$ | $\beta$ | $P$ |
| cg16672562 | -0.071 | 0.299 | -0.084 | 0.220 | -0.027 | 0.682 | -0.062 | 0.369 |
| cg22891070 | -0.002 | 0.974 | -0.060 | 0.374 | 0.004 | 0.946 | 0.021 | 0.755 |
| cg27146050 | -0.085 | 0.215 | -0.161 | 0.029 | -0.095 | 0.156 | -0.095 | 0.165 |
| mean | -0.059 | 0.400 | -0.104 | 0.140 | -0.039 | 0.569 | -0.047 | 0.505 |

Adjusted for age, systolic blood pressure, hemoglobin A1c, %neutrophil, smoking habit and exercise habit. $P < 0.05$ was considered statistically significant.

Smoking habits alter the status of DNA methylation^27, 28^. Therefore, we examined whether DNA methylation of the different regions of *HIF3A* were associated with the parameters of obesity in non-smokers (i.e., the participants excluding current smokers and non-smokers who stopped smoking within the last 5 years; Table 6). There was no significant correlation between DNA methylation level at each CpG site and the parameters of obesity in men. In women, there were significant correlations between DNA methylation level at cg27146050 and BMI, VAT thickness, and % body fat (*P*<0.05).

**Table 6.** Multiple linear regression analysis for correlations between HIF3A gene DNA methylation and obesity parameters in non-smokers.

| **Men** |  |  |  |  |  |  |  |  |
| --- | --- | --- | --- | --- | --- | --- | --- | --- |
| **CpG site** | **BMI** |  | **VAT** |  | **SAT** |  | **% Body fat** |  |
| | $\beta$ | $P$ | $\beta$ | $P$ | $\beta$ | $P$ | $\beta$ | $P$ |
| cg16672562 | 0.086 | 0.306 | 0.098 | 0.228 | -0.046 | 0.581 | 0.068 | 0.426 |
| cg22891070 | 0.101 | 0.233 | 0.040 | 0.627 | 0.004 | 0.963 | 0.030 | 0.727 |
| cg27146050 | -0.017 | 0.847 | 0.012 | 0.883 | -0.027 | 0.754 | -0.067 | 0.449 |
| mean | 0.077 | 0.368 | 0.059 | 0.475 | -0.228 | 0.790 | 0.020 | 0.821 |
| **Women** |  |  |  |  |  |  |  |  |
|  | **BMI** |  | **VAT** |  | **SAT** |  | **% Body fat** |  |

| CpG site | $\beta$ | $P$ | $\beta$ | $P$ | $\beta$ | $P$ | $\beta$ | $P$ |
| --- | --- | --- | --- | --- | --- | --- | --- | --- |
| cg16672562 | -0.009 | 0.103 | -0.075 | 0.311 | -0.050 | 0.492 | -0.110 | 0.133 |
| cg22891070 | -0.059 | 0.413 | -0.068 | 0.345 | -0.007 | 0.924 | -0.019 | 0.791 |
| cg27146050 | -0.152 | 0.040 | -0.179 | 0.016 | -0.110 | 0.132 | -0.162 | 0.029 |
| mean | -0.124 | 0.096 | -0.114 | 0.129 | -0.056 | 0.449 | -0.102 | 0.170 |

Adjusted for age, systolic blood pressure, hemoglobin A1c, %neutrophil and exercise habit, and excluded for current smoker (include stopped smoking less than 5 years). $P < 0.05$ was considered statistically significant.

## Discussion

We determined the association between DNA methylation at three CpG sites (cg16672562, cg22891070, and cg27146050) of the intron 1 of *HIF3A* and parameters of obesity in the general Japanese population. There was a significant difference in the DNA methylation of *HIF3A* between the sexes. Multiple linear regression analysis showed a correlation between DNA methylation at cg27146050 in *HIF3A* and thickness of VAT in women. Excluding current smokers and non-smokers who stopped smoking within the last 5 years, there correlations between DNA methylation at cg27146050 in *HIF3A* and thickness of VAT thickness, BMI, and % body fat in women.

In this study, DNA methylation at each CpG site of intron 1 of *HIF3A* was higher in women than those in men. This was consistent with that reported by Main et al.^21^. This difference between men and women suggests that women exhibit lower expression of *HIF3A* during hypoxia than the expression in men under similar conditions owing to differential capacities of gene regulation. Women exhibit a relatively less pronounced physiological response to hypoxic stress than that in men^32^. This can be attributed to the increase in DNA methylation of *HIF3A* in women.

Dietary factors, such as nutrition, cause a change in DNA methylation^12^. We have recently demonstrated that the intake of dietary vitamin affects lipid profiles via the modulation of DNA methylation within lipid-related genes^16^. DNA methylation variants of *HIF3A* are associated with alterations in BMI based on the consumption of total vitamins or supplemental vitamin B^33^. Therefore, it is possible that the intake of vitamins or other nutrients causes a change in DNA methylation in *HIF3A*, thereby resulting in the development of obesity.

Tobacco smoking is another environmental factor that affects the incidence of obesity^34^. DNA methylation positively correlates with smoking habits^27, 28^. Thus, smoking habits may influence the association between *HIF3A* DNA methylation and the parameters of obesity. We determined whether DNA methylation in *HIF3A* associated with parameters of obesity in non-smokers (excluding current smokers and non-smokers who stopped smoking within the last 5 years). The correlation between DNA methylation at various sites of *HIF3A* and parameters of obesity increased in this population than that including non-smokers and smokers. To the best of our knowledge, this is the first report on the correlation between DNA methylation levels at the CpG sites in *HIF3A* and parameters of obesity, such as thickness of VAT and smoking habits, in the general Japanese population. However, smokers were not excluded from the group of non-smokers. Therefore, future studies should focus on determining the association between DNA methylation in *HIF3A* and parameters of obesity within non-smokers.

Dick et al.^20^ demonstrated the correlation between DNA methylation levels at three CpG sites (cg16672562, cg22891070, and cg27146050) in intron 1 of *HIF3A* in the blood and BMI; this has also been confirmed by other studies^21, 34^. In women, DNA methylation at cg27146050 correlated with the thickness of VAT (based on multiple linear regression analysis). However, this association has been reported in men^20^. This discrepancy may be explained attributed to the differences in DNA methylation of *HIF3A* and smoking habits of men and women. Thus, future studies are warranted to elucidate the extent of DNA methylation in *HIF3A* between men and women.

Dick et al.^20^ used a microarray to demonstrate the association between *HIF3A* DNA methylation and BMI in humans. Microarrays are useful in understanding global DNA methylation. However, this technique cannot measure methylation using immobilized methylated probes and exhibits poor quantification. Thus, this study employed pyrosequencing to analyze DNA methylation in *HIF3A*. This method is excellent in quantifying the extent of DNA methylation at selected CpG sites in specific target genes. Thus, pyrosequencing provides a more reliable scenario of the association between DNA methylation at CpG sites of intron 1 of *HIF3A* and parameters of obesity in the general Japanese population than the correlation reported by Dick et al.^20^.

Finally, Hatanaka et al.^35^ showed that the ectopic expression of *HIF3A* induces the expression of several adiposity-associated genes in 3T3-L1 cells. This suggests that low levels of DNA methylation in *HIF3A* upregulates *HIF3A*, thereby resulting in adiposity. Accordingly, we observed a positive correlation between DNA methylation in *HIF3A* and thickness of VAT (that directly reflects obesity as compared to BMI) in women. Therefore, the thickness of VAT has important clinical implications in obesity-related diseases.

Taken together, the DNA methylation level at cg27146050 of intron 1 of *HIF3A* correlated well with parameters of obesity in non-smokers of the general Japanese women. This study has some limitations. First, the data do not show a causal relationship between DNA methylation at different sites of *HIF3A* and parameters of obesity since this was a cross-sectional study. Second, this study analyzed a small sample size. Finally, we did not determine alterations in the mRNA levels of *HIF3A*.Thus, future studies should focus on analyzing the association between DNA methylation level in *HIF3A* and the parameters of obesity over a longer period using a larger sample size. Furthermore, we will attempt to analyze the mRNA and protein levels of *HIF3A* in the blood of participants.

## Conclusion

This is the first study to report the correlation between DNA methylation at CpG site in *HIF3A* and parameters of obesity, such as thickness of visceral adipose tissue and smoking habit, in the general Japanese population. DNA methylation of the CpG sites of *HIF3A* may be associated with body mass index.

## Acknowledgments

We thank the participants and staff of the Health Examination Program for Residents of Yakumo, Hokkaido, Japan.

